# Deciphering the Nrf2/ARE Mechanism: GanCaoXieXin Decoction Combats Oxidative Stress in Ulcerative Colitis Pathogenesis

**DOI:** 10.1101/2025.07.02.662699

**Authors:** Ping Ling, Zhang Bing, XinRui Zhang, Yingchao Liu, Yan Shen

## Abstract

**Background:** GanCaoXieXin (GCXX) decoction, a classic prescription, has shown clinical efficacy in treating ulcerative colitis (UC). However, its mechanism remains incompletely understood.

**Objective:** This study aims to explore how GCXX modulates the Nrf2/ARE signaling pathway to mitigate oxidative stress (OS)-induced damage and thereby ameliorate UC.

**Materials and methods:** Network pharmacology and bioinformatics analyses identified key targets of GCXX in UC treatment. Ultra-performance liquid chromatography combined with quadrupole time-of-flight mass spectrometry (UHPLC-MS/MS) analyzed GCXX’s effective compounds. A 2.5% Dextran Sulfate Sodium Salt (DSS)-induced UC mouse model was used. Immunohistochemistry (IHC) assessed tight junction proteins. Enzyme-linked immunosorbent assay (ELISA) measured intestinal permeability and oxidative stress markers. Western blot (WB) analyzed Nrf2/ARE signaling proteins. In 800μmol/L H2O2-induced oxidative stress (OS) state HT-29 cells, cell viability, apoptosis, oxidative stress indicators, and apoptosis-related proteins were evaluated. Immunofluorescence (IF) detected Nrf2/ARE signaling axis proteins.

**Results:** The results of network pharmacology and bioinformatics analysis revealed that GCXX could intervene in UC by regulating OS-related pathways. GCXX contained antioxidant components like quercetin, berberine, and baicalin. *In vivo*, GCXX alleviated mucosal damage, reduced intestinal permeability, downregulated MDA, upregulated SOD, suppressed Keap1, and promoted tight junction and Nrf2/ARE pathway proteins. *In vitro*, GCXX increased cell survival, improved antioxidant capacity, reduced MDA and apoptosis in OS state cells. Immunofluorescence confirmed Nrf2/ARE pathway as crucial in GCXX’s protective effects.

**Conclusions:** GCXX elevates the expression level of Nrf2, HO-1 and NQO1, thus reducing entercell OS-induced damage, intestinal cell apoptosis, alleviating UC intestinal mucosal damaget.

## 1. Introduction

Ulcerative colitis (UC), an etiologically unclear chronic inflammatory bowel illness ^[(1)]^, features multiple superficial ulcers in the affected colon mucosa, accompanied by congestion, edema, and clinical manifestations of abdominal pain, hematochezia, and chronic diarrhea ^[(2)]^. Its pathogenesis remained unraveled, but it is postulated to arise in genetically predisposed individuals following environmental exposure, and be closely related to intestinal epithelial barrier damage, intestinal flora and dysimmune response ^[(3–5)]^. A global epidemiological study on inflammatory bowel disease (IBD) indicates that while the occurence and prevalence of IBD in East Asia are currently lower than those in Western countries, there is a notable upward trend, with an increasing number of new cases over time ^[(6)]^. Recent years have witnessed the rapid rise in the incidence rate and prevalence of UC in China with a growing number of critical cases, predominantly affecting a younger population, thereby significantly impacting social productivity and individual life^[(7)]^. GCXX, a traditional Chinese medicine(TCM) prescription, has exhibited therapeutic efficacy in the UC treatment, and can effectively relieve colitis in the UC model mice and protect the intestinal mucosal barrier^[(8)]^. Nevertheless, relevant mechanism remains obscure due to multiple componenst, targets and pathways. Therefore, further exploration is necessary.

Oxidative stress (OS) refers to a pathophysiological state featuring disruption of cell’s normal aerobic energy metabolism process and disturbance of redox state^[(9)]^. Chronic inflammation, pathogen infection, ischemia and reperfusion injury, can lead to excessive ROS production^[(10, 11)]^. To maintain homeostasis, the body initiates a clearance program when it senses substances like ROS. However, OS-induced damage occurs when the organism’s own clearance capacity is insufficient to eliminate the overproduced ROS. Studies have proved that OS is a critical pathological mechanism causing inflammation, necrosis and ulcer formation of UC colon tissue, and also plays a central role in itsdeterioration ^[(12, 13)]^. Therefore, the regulation of OS levels induced by intestinal inflammation is essential for mitigating intestinal mucosal injury in UC.

This study utilizes advanced network pharmacology^[(14)]^ and bioinformatics analyses to elucidate the mechanisms by which GCXX ameliorates UC. Ultra-performance liquid chromatography combined with quadrupole time-of-flight mass spectrometry (UPLC-Q-TOF-MS) was used to investigate the active ingredients in GCXX. Dextran Sulfate Sodium Salt (DSS) helped to construct UC mice model^[(15)]^, and therapeutic efficacy was assessed by monitoring parameters such as disease activity index (DAI), body weight, colon macroscopic damage index (CMDI), colon length and tissue damage index (TDI). By detecting indicators such as MDA and SOD, we assessed the ameliorative influence of GCXX on OS-led damage in UC. Moreover, an OS state *in vitro* intestinal cell model assisted in investigating the effect of GCXX on OS-induced damage and apoptosis level in OS fine cell model. We clarified that GCXX protected the intestinal barrier by downregulating OS-induced damage and apoptosis, and further elucidated that the protective effect of GCXX is mediated through Nrf2/ARE signaling axis. Therefore, our study seeks to validate that regulation of Nrf2/ARE signaling axis by GCXX attenuates enterocyte OS-induced damage and apoptosis, repairs the intestinal barrier, and ameliorates UC.

## 2. Materials and Methods

### 2.1. Reagents and antibodies

The reagents and antibodies used in this study are shown in the Table S1 and 2.

### 2.2 Preparation of GCXX

The GCXX formulation consists of 12 g of fried licorice, 9 g of dried ginger, pinellia and Scutellaria baicalensis, 9 g of Dangshen, 6 g of jujube,and 3 g of Coptis chinensis. Procured from Hangzhou Binjiang Chinese Medicine Clinic of Zhejiang Chinese Mediccal University (Hangzhou, Zhejiang, China), all ingredients were certified by pharmacologists. The formulation was then concentrated to a liquid form with a density of 1 g/mL using a rotary evaporator and stored at −20℃.

### 2.3 Drug-containing serum preparation

Based on pharmacological guidelines for dose conversion between humans and rats, the equivalent dose of clinical GCXX in rats was 6 g/kg/day. Twenty specific pathogen-free (SPF) male Sprague-Dawley (SD) rats were randomly assigned into two groups: one received GCXX (6 mL/kg/day), and the control had the same volume of distilled water twice daily at 12-hour intervals for three days. On day 4, one hour after the last gavage (12 hours post-administration), we collected blood from the abdominal aorta. Serum was isolated under sterile conditions, and inactivated in 56℃ constant temperature water bath for 30 minutes, which was followed by 0.22μm microporous membrane filtration and storage at −80℃.

### 2.4 Network pharmacological analysis of the GCXX-UC

All active compounds in GCXX were searched with the use of Traditional Chinese Medicine Systems Pharmacology Database and Analysis Platform (TCMSP). Compounds that meet the threshold of OB>=30%, DL>=0.18, and HL>=4H were identified as effective. PubChem and SwissTargetPrediction were used to search the prediction target. UC-related targets were filtered from the GeneCards and OMIM databases by using the keyword “ulcerative colitis”. A Venn diagram assisted in identifying the intersection targets of TCM active compounds and UC-related targets, indicating potential targets in UC.

Subsequently, these candidate targets were entered into the STRING database for protein-protein interaction (PPI) analysis and also into R version 4.3.2 for additional evaluation. We performed Gene Ontology (GO) and Kyoto Encyclopedia of Genes and Genomes (KEGG) functional enrichment analysis of genes with visualized results.

### 2.5 Bioinformatics analysis of the GCXX

The UC dataset GSE243625, comprising 15 UC and normal samples respectively, was downloaded from GEO for differential gene expression analysis for detecting the differentially expressed genes (DEGs). Disease-related modules were obtained via the WGCNA package of R, and correlation matrices between module eigengenes were computed. Differential genes, hub genes of WGCNA, and GCXX target genes were selected with the help of the clusterProfiler package (v 4.6.2), and the org.Hs.eg. db package (v 3.16.0) for GO and KEGG enrichment analysis (p <0.05 and q < 0.05). GOplot was utilized to visualize the results..

### 2.6 UPLC-Q-TOF-MS analysis of GCXX

200 μ L of GCXX was refined and diluted to 100 mg/mL with methanol and centrifuged at 12,000 g for 10 minutes. The supernatant was used for UH PLC-Q-TOF-MS analysis (Waters, Ireland). Chromatographic conditions were: ACQUITY UPLC HSS T3 column (100 mm 2.1 mm, 1.8 µ m; Waters Corporation, Ireland), flow rate of 0.3 mL/min, sample volume of 2.00 μ L, sampler temperature of 4℃, and column temperature of 40℃. Mobile phases A and B were 0.1% (v / v) formic acid / water and 0.1% (v / v) formic acid / acetonitrile. Elution gradient was: 0 - 2min (5% B), 2 - 10min (15% B), 10-15 min (25% B), 15 - 18min (50% B), 18 - 23min (100% B), 23 - 25min (2% B), 25 - 30min (2% B); The column temperature was 40℃, and the injection volume was 2.00 μ L. The mass spectrometric analysis was performed under the following conditions: in positive ion mode, capillary voltage at 3.5 kV, sampling cone voltage at 40.0 V, source temperature:150℃, desolvation temperature: 350℃, cone gas flow: 80℃, desolubilized gas flow: 800.0 L/h, scanning time: 0.10s, interval time: 0.014s, and mass range: 50-2000m/z. In negative ion mode, the parameters were: capillary voltage at 3.0 kV, sampling cone voltage at 40.0 V, source temperature: 120℃, desoluble temperature: 360℃, cone gas flow: 50℃, desoluble gas flow: 800.0 L/h, scanning time: 0.10s, scanning time: 0.014s, and mass range: 50-2000 m / z. All data were acquired with the help of MassLynx V4.1 (Waters Corporation, Milford, MA, USA) and subsequently imported into Progenesis QI V2.0 (Waters Corporation, Milford, MA, USA) for pre-processing.

### 2.7 Experimental animals and the cells

We procured SPF male C57BL / 6 mice (6-8 weeks, 20-22 g) from Hangzhou Qizhen Laboratory Animal Technology Co., Ltd. (SCXK (Zhejiang) 2022-0005, Hangzhou, China). Maintained at the Animal Laboratory Center of Zhejiang Chinese Medical University (ZCMU) at 22 ± 2℃ with a 12-hour light/dark cycle, they could have standard food and sterilized water freely. All animal experiments were approved by the Experimental Animal Ethics Committee of ZCMU (IACUC-20231120-28) and conducted at ZCMU. Animal welfare work was conducted as per the Guidelines on the Management and Use of Laboratory Animals (Ministry of Science and Technology, China, 2016).

Human colon cancer cells (HT-29) were bought from Mirror Qidian (Shanghai) Cell Technology Co., Ltd. (Cellverse Bioscience Technology Co., Ltd.).

### 2.8 Preparation and group administration of animal models

After a one-week acclimatization period, random allocation was carried out to assign the animals into six groups: 5-ASA group, Low-GCXX group, Middle-GCXX group, High-GCXX group, UC group and NC group (n=10). The UC, 5-ASA and three dose GCXX groups consumed drinking water with 2.5% DSS for a week ^[(16)]^, and the NC group drank distilled water freely. Based on pharmacological guidelines for dose conversion between humans and rats, the equivalent dose of GCXX in mice was 8.55 g/kg/day. This concentration was set as the high dose, with medium and low doses established by equivalent-fold serial dilution. Following the setablishment of UC mouse model, the high-dose GCXX group received GCXX (8.55 mL/kg/day), while the medium- and low-dose groups were administered 4.275 mL/kg and 2.1375 mL/kg, respectively, oral gavage for 7 days, the 5-ASA group received 5-aminosalicylic acid (150 mg/kg/day) via oral gavage for 7 consecutive days, and NC and UC groups were given the same volume of distilled water.

### 2.9 DAI score

Throughout the experiment, the general condition and stool characteristics were observed, and the body weight was measured every two days. DAI was scored once every two days according to stool viscosity, the presence of stool bleeding and the proportion of mouse body weight loss as detailed in Table S3.

### 2.10 Tissue collection and CMDI score

After the end of 7-days treatment, blood was collected from the posterior venous plexus of mouse eyeball, and centrifuged at 3000 rpm for 10 minutes. We collected and stored the serum at −80℃. Mice were euthanized with CO_2_, the colon was removed, and the lesions of the colon were observed for CMDI scoring (Table S4), and the colon length was measured and photographed. 1cm intestinal segments were fixed in 4% paraformaldehyde, and the rest was stored at **-**80℃.

### 2.11 Histopathological test (HE) and tissue TDI score

Colons fixed in 4% paraformaldehyde were embedded in paraffin, sectioned, stained with hematoxylin and eosin (HE), and observed via a light microscope. TDI scores were determined based on epithelial mucosal injury and inflammatory infiltration as described in Table S5.

### 2.12 Immunohistochemistry (IHC)

The distribution characteristics of Nrf2, Keap1, HO-1, and NQO1,Occludin, Claudin-1, and ZO-1 were observed and measured by IHC. IHC staining was semi-quantitatively analyzed using Image J software.

### 2.13 Enzyme-linked immunosorbent assay (ELISA)

After the sample processing, OD was measured at a wavelength of 450nm. The standard curve, with the concentration of the standard on the x-axis and the OD value on the y-axis, was used to determine the sample concentrations, which were then used to calculate the DAO and D-LAC levels.

### 2.14 Cell culture and group administration

HT 29 cells were seeded in 50 mL culture flasks containing McCoy’s medium with 10% inactivated FBS and 1% streptomycin, and incubated at 37°C in 5% CO_2_ with saturated humidity.

All cells were divided into NC, OS, low dose (Low-GCXX), high dose (High-GCXX) GCXX,si-NC + OS + GCXX, and si-Nrf2 + OS + GCXX groups. The normal control group was cultured in a complete medium for two days; the OS group was pre-cultured in a complete medium for one day, and replaced with 800 μmol / LH_2_O_2_ ^[(17)]^ for 24 hours; Low-GCXX and High-GCXX groups were pre-cultured in medium with 10% and 20% GCXX containing serum for 24 hours, and replaced with 800μmol / L H_2_O_2_ for 24 hours; The si-NC + OS + GCXX group and si-Nrf2 + OS + GCXX group, after transfection with NC siRNA and Nrf2 siRNA, were cultured in 20% GCXX serum and then treated with medium containing 800 μmol / L H_2_O_2_ respectively for 24 hours.

### 2.15 MTT

After the cells were administration as described above, 20 μL MTT solution (5g / L) was put into all wells at 37℃ for 4 hours. Following centrifugation, MTT solution was aspirated, 150 μ L dimethylsulfoxide was added to each well. The absorbance (A) at 490nm was calculated through a microplate reader. Finally, the viability of cells was calculated. Each group was repeated thrice.

### 2.16 Flow Cytometry

After dosing as above, cells were collected by centrifugation. We added 5 μL Annexin V-APC and 10 μL 7-ADD to the suspension. The mixture was set with single dye tubes, gently mixed, and incubated for 5 minutes at room temperature. Apoptosis was detected by analyzing the cells: Annexin V-APC was used for the FL4 channel, and 7-ADD was used for the FL3 channel. Each experiment was repeated three times.

### 2.17 Small interfering RNA (siRNA) transfection

2 μg of plasmid and 10 μL liposomes was added to 200 μL basal medium at room temperature for 5 minutes; was added to 200 μL basal medium; The plasmid and liposomes were mixed and incubated at room temperature for hal of an hour to prepare transfection complexes, which were then added to cells for transfection. The medium was changed after 4-6 hours. The Nrf2 siRNA and its negative control (si-NC) were constructed and introduced into HT 29 cells through the aforementioned lipofection method.

### 2.18 Immunofluorescence (IF)

After administration, cells were washed with PBS, and fixed with 4% paraformaldehyde before permeabilization with 0.1% Triton X-100 for 20 minutes. Blocking was performed with 5% BSA for half of an hour. They were incubated overnight at 4℃ with primary antibodies against Nrf2, HO-1, and NQO1. Cells were then washed with PBS and incubated for 1-2 hours with secondary antibodies. After washing, nuclei were stained with 10 ng/mL DAPI for 15 minutes in the dark at 4℃. Each experiment was repeated thrice.

### 2.19 Western blotting (WB)

After 30 μg of protein from each group for sodium dodecyl sulfate polyacrylamide gel electrophoresis (SDS-PAGE), the proteins were transferred to polyvinylidene fluoride (PVDF) membrane. Primary antibody against target protein was added and incubated overnight at 4℃. After the membrane was washed, the secondary antibody was added and incubated for 1 hour at room temperature.1 After wash in TBST, the film was developed with ECL chemiluminescence. Image processing software helped to analyze the band gray values. GAPDH was used for internal reference. All experiments was performed three times.

### 2.20 Measurements of OS indexes such as MDA, SOD and T-AOC

MDA, SOD and T-AOC were measured through kit biochemistry and operated according to instructions for the kit. Each experiment was repeated thrice.

### 2.21 Statistical analysis

Statistical software SPSS 29 was employed for data analysis and processing. Data are presented as mean ± standard deviation. The Shapiro-Wilk test assisted in the assessment of data distribution. The analysis of variance statistical test helped with data comparison. When the data were not normally distributed, we employed the Kruskal-Wallis test. Statistical significance was defined as a P-value less than 0.05..

## 3. Results

### 3.1 Network pharmacology and bioinformatics results predicted GCXX’s Intervention in UC by regulating OS-related pathways

In the GeneCards and OMIM databases, we identified UC-related targets using the term "ulcerative colitis", and 5,828 genes with potential targets were obtained after duplicates were removed. A Venn diagram comparing these genes with 595 genes corresponding to 343 active components in GCXX, yielded 347 potential targets for GCXX treatment in UC (Fig. 1A). 347 potential targets were introduced into STRING to elucidate PPI, which were subsequently visualized via Cytoscape 3.8.0 (Fig. 1B). The top PPI results included non-receptor tyrosine kinase (SRC), phosphatidylinositol 3-kinase (PIK3R1), mitogen-activated protein kinase 3 (MAPK3). The 347 genes were imported into R 4.3.0 for GO / KEGG functional enrichment analysis as well as visualization of the results (*P*<0.05). GO functional enrichment analysis yielded 3,975 GO entries, including 3,185 BP, 270 CC and 520 MF entries, and top 10 entries were selected to draw bars (Fig. 1C). Among them, BP mainly involved reactions to chemicals, reactions to oxygen compounds and response to stimuli; CC mainly involved cell periphery, plasma membrane and growth process compartment; MF mainly involved protein kinase activity, kinase activity and transfer of phosphorus-containing groups. Additionally, the KEGG enrichment analysis yielded 183 signaling pathways, with the top 20 by P-value primarily involving chemical carcinogenic—reactive oxygen species, PI3K-Akt, and MAPK signaling pathways (Figure. 1D). GO and KEGG results demonstrated functional annotation and signaling pathways related to OS, indicating that GCXX can interfere with UC by modulating OS-related mechanisms. UC dataset GSE243625 was selected from GEO database for differential gene analysis, and a total of 3,685 differential genes were identified, with 1,578 genes downregulated and 2,107 upregulated. The volcano map and heat map of the differential genes were drawn (Fig. 2A and B). The disease-related module was identified by the WGCNA package in R (Fig. 2C-E). We selected the black, brown, tan modules, with hub gene threshold of GS> 0.6 and MM> 0.6. Specifically, the black, brown and tan modules contained 261, 857 and 1,223 genes, respectively. We then calculated the correlation coefficient matrix between the module feature vectors and generated a heatmap to visualize the correlation (Fig. 2F-H). 243 differential genes, WGCNA hub genes, and GCXX target genes (Fig. 2I-K) were subjected to GO and KEGG enrichment analysis (Fig. 2L and M). GO functional enrichment analysis yielded 2.437 GO entries, including 2,219 BP, 97 CC and 121 MF entries, and top 10 entries were selected to draw the bar graph. Among them, BP mainly involved active regulation of cytokine production, cell response to biological stimuli and damage repair; CC mainly involved membrane raft, membrane microdomain and basolateral plasma membrane; MF mainly involved the activity of oxidoreductase, protein heterodimerization activity and cytokine receptor binding. KEGG enrichment analysis yielded 111 items, including 11 related to cellular processes, 11 to environment information processing, 54 to human diseases, 11 to metabolism, and 24 to organismal systems, with top 5 pathways selected for visualization. KEGG mainly involved PI3K, Akt, NF-kappaB, and TNF signaling pathways. Results of the bioinformatics analysis revealed an enrichment of the molecular functions associated with OS, suggesting that the regulation of OS pathway is most likely the mechanism through which GCXX ameliorates UC.

**Figure 1.**
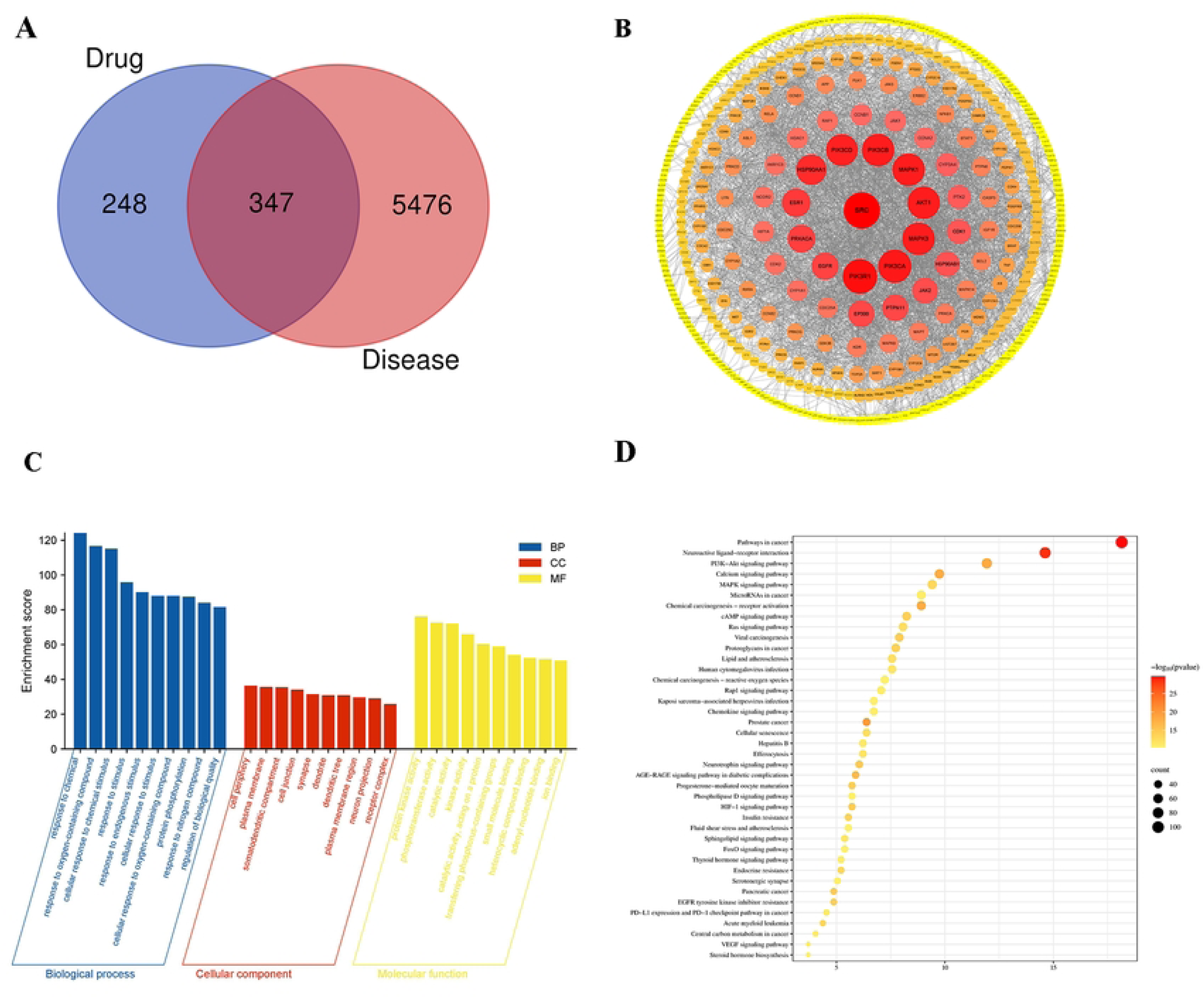
Results of the pharmacological analysis of the GCXX-UC network. (A) Wayn diagram of target intersection between GCXX and UC. (B) PPI network analysis map of key targets. (C) GO enrichment analysis, showing top 10 items. (D) KEGG analysis, showing top 40 items.

**Figure 2.**
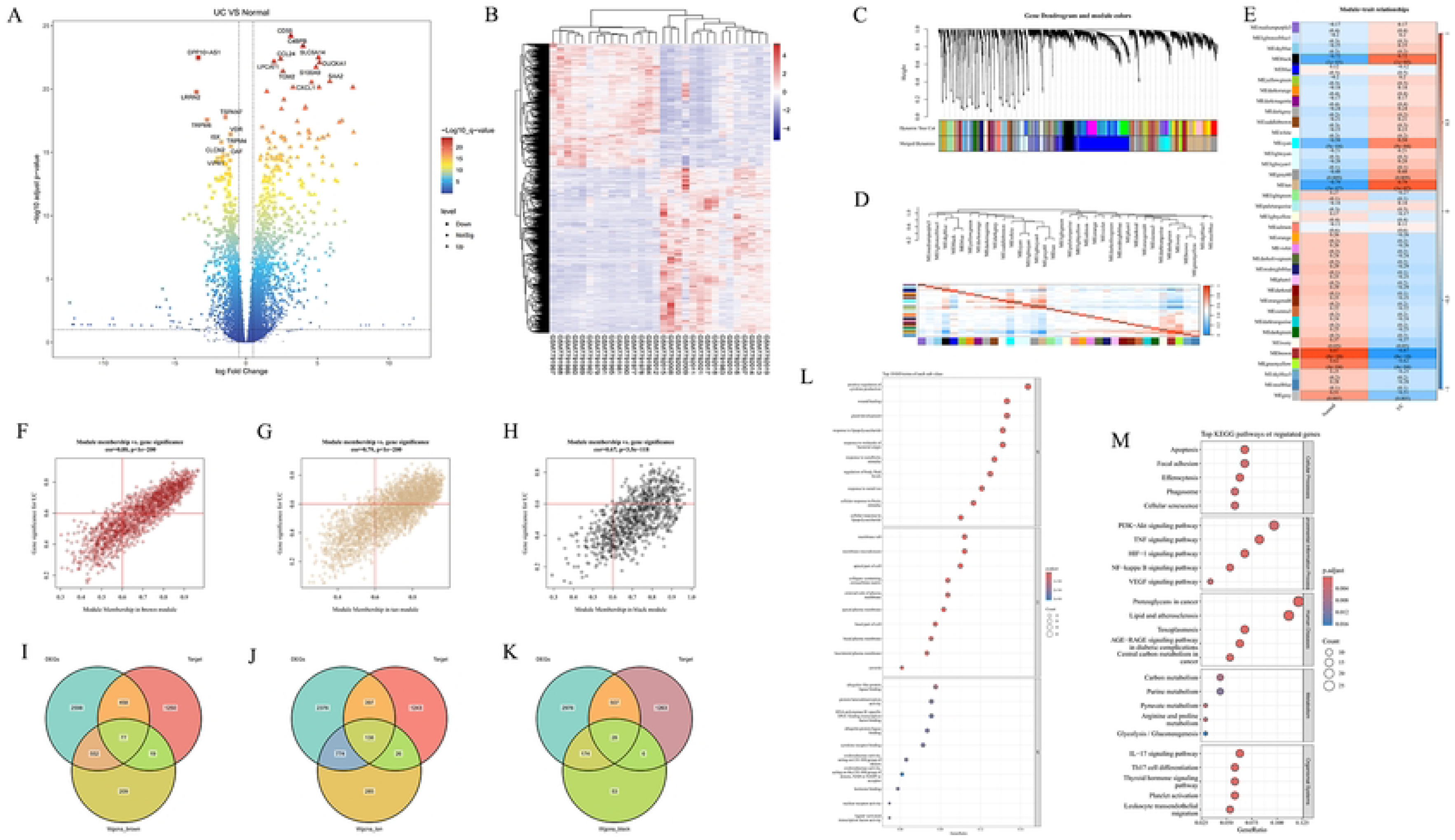
UC differential gene target screening and enrichment analysis. (A-B) Volcano map and heat map of UC differential genes. (C-E) WGCNA hub module screening. (F-H) Extraction of the key modules. (I-K) Differential genes, hub genes of WGCNA, intersection targets of drugs. (L-M) GO and KEGG enrichment analysis of intersection targets.

### 3.2 GCXX high-performance liquid phase mass spectrometry results

UHPLC-Q-TOF-MS detected 20,523 compounds in GCXX (Fig. 3A and B), including 10 major compounds such as quercetin, berberine and baicalin (Table 1), and molecular formula can be found in Fig. 3C. GCXX detected many monomer components with antioxidant activity, such as quercetin, berberine, and baicalin.

**Figure 3.**
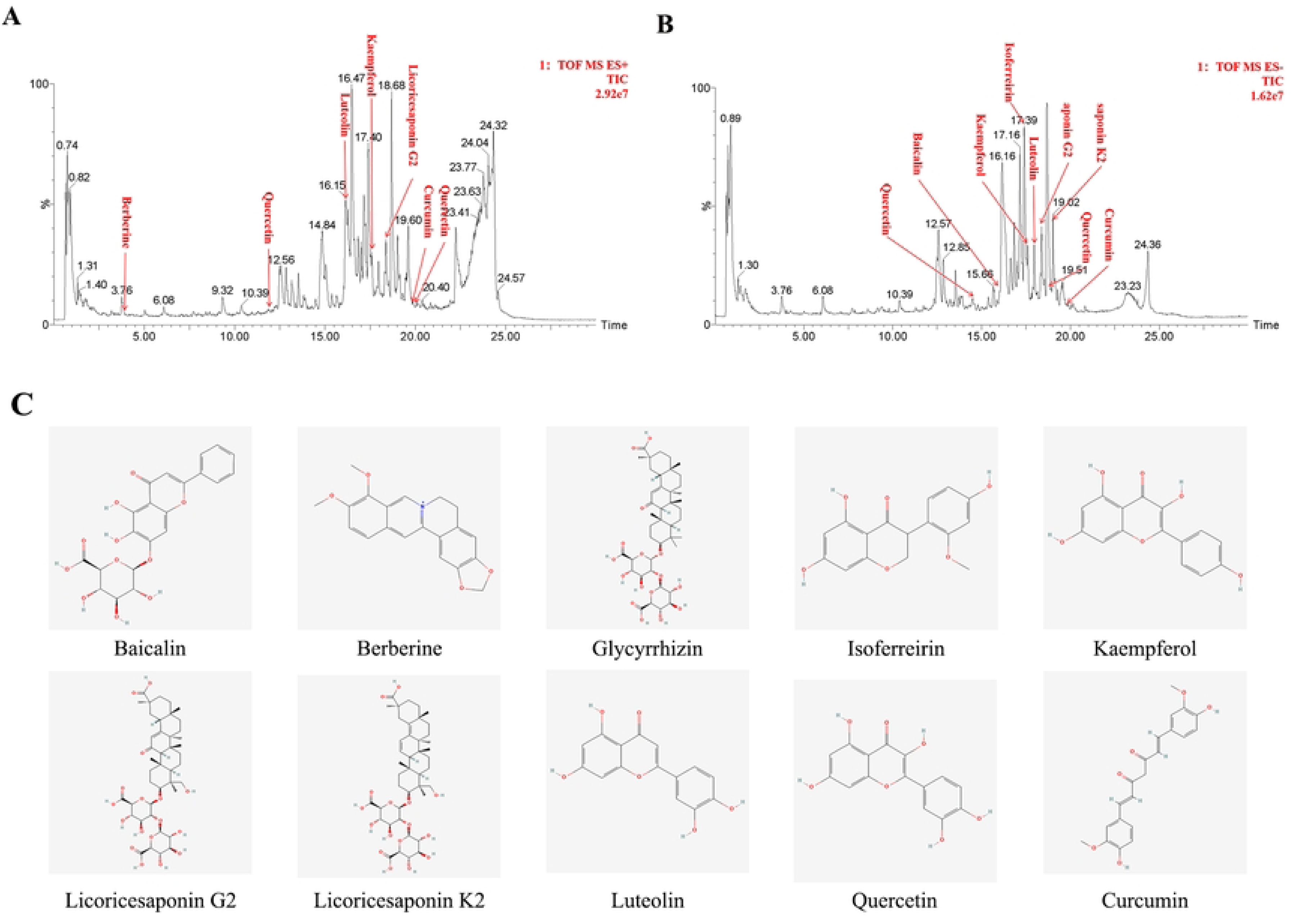
Results of the GCXX UHPLC-Q-TOF-MS analysis. (A) TIC profile in positive ion mode. (B) TIC profile in negative ion mode. (C) Structures of 10 major compounds.

**Table 1.**
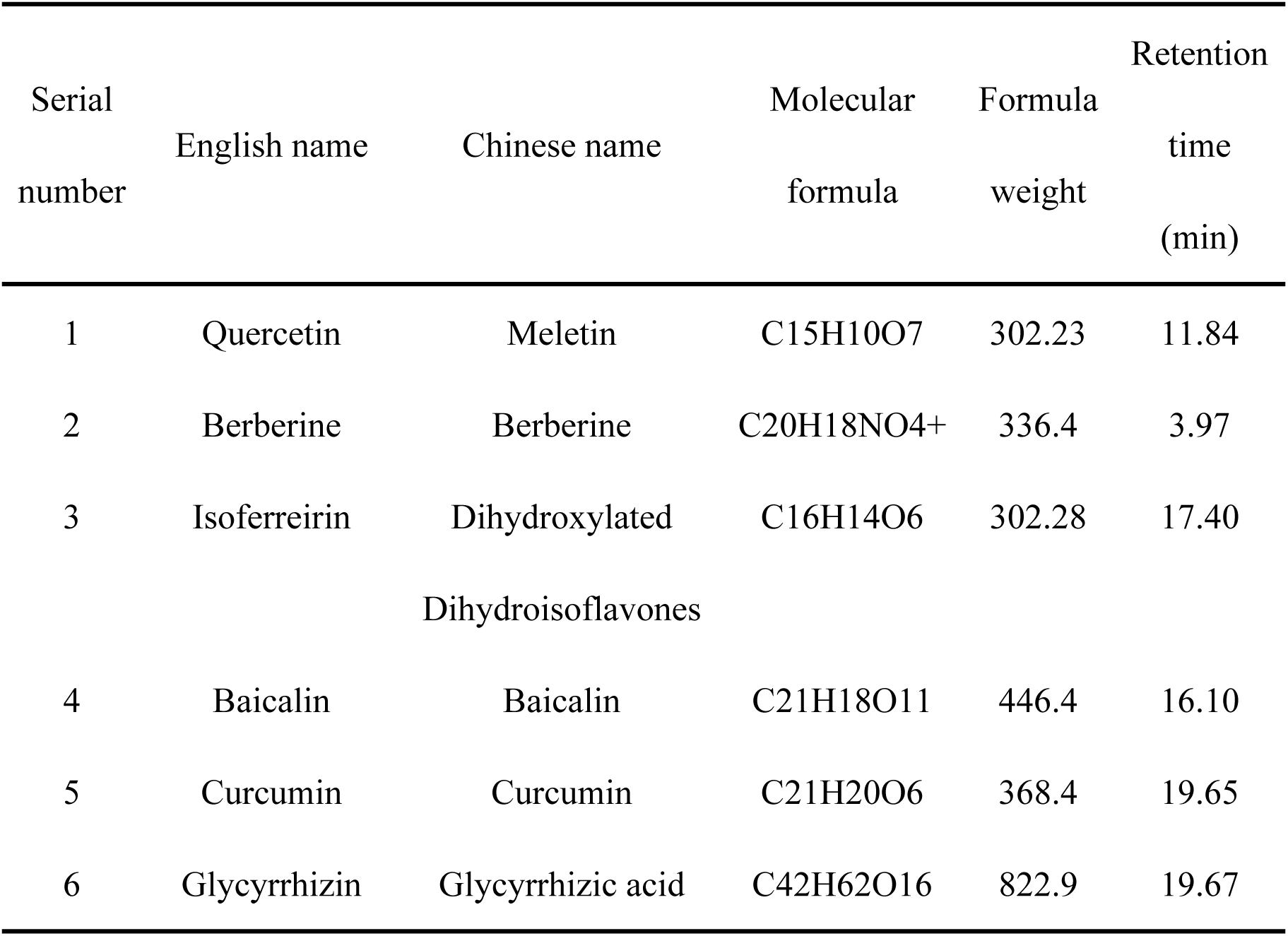
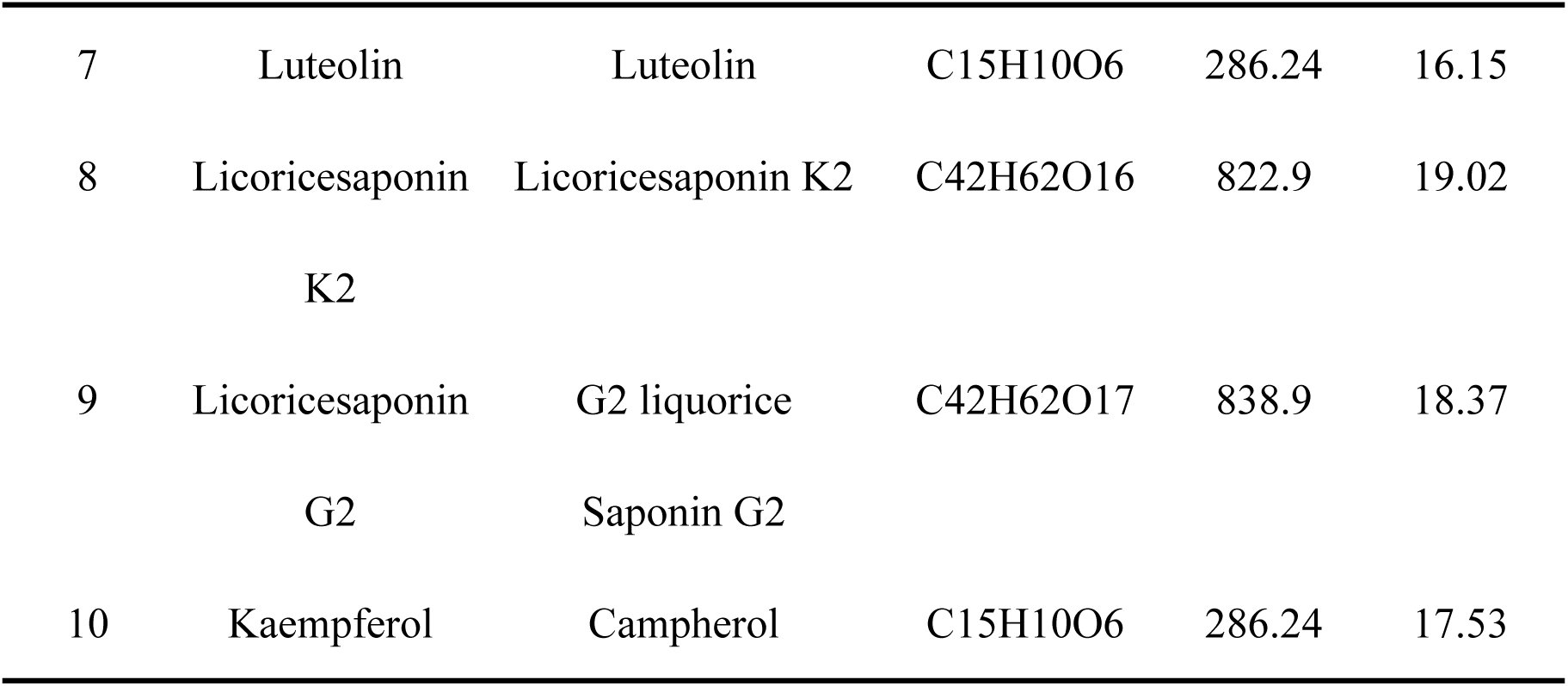
10 major compounds in GCXX.

### 3.3 GCXX attenuated colitis in DSS-induced UC mice

Markedly lowered viability and body weight were observed in the UC group in reference to the NC group, whereas low, medium and high-dose GCXX groups demonstrated an improvement in both viability and body weight in relation to the UC group, with the high-dose group exhibiting the most significant recovery. Prior to death, the body weight of the UC group was notably lower than the control, while low, medium and high-dose GCXX groups showed a slightly elevated body weight in comparison to the UC group (Fig. 4A) DAI score was assessed every two days following the establishment of DSS-induced UC. Following GCXX administration, a gradual decrease of DAI score in the low, medium and high-dose GCXX groups was observed, and upon the end of administration, the DAI score was notably lower than that in the UC group (Fig. 4B). Colon length was measured after mice death on day 14. As shown in Figure, the UC group had a markedly shorter colon length in reference to the control, while low, medium and high-dose GCXX groups presented with a recovery in colon length compared to UC group (Fig. 4C). In addition, the colon mucosal injury was evaluated after killing, and CMDI score was performed (Fig. 4D). The CMDI score for the UC group was substantially higher than that of the control, whereas the score for the experimental group was markedly lower in reference to the UC group.

**Figure 4.**
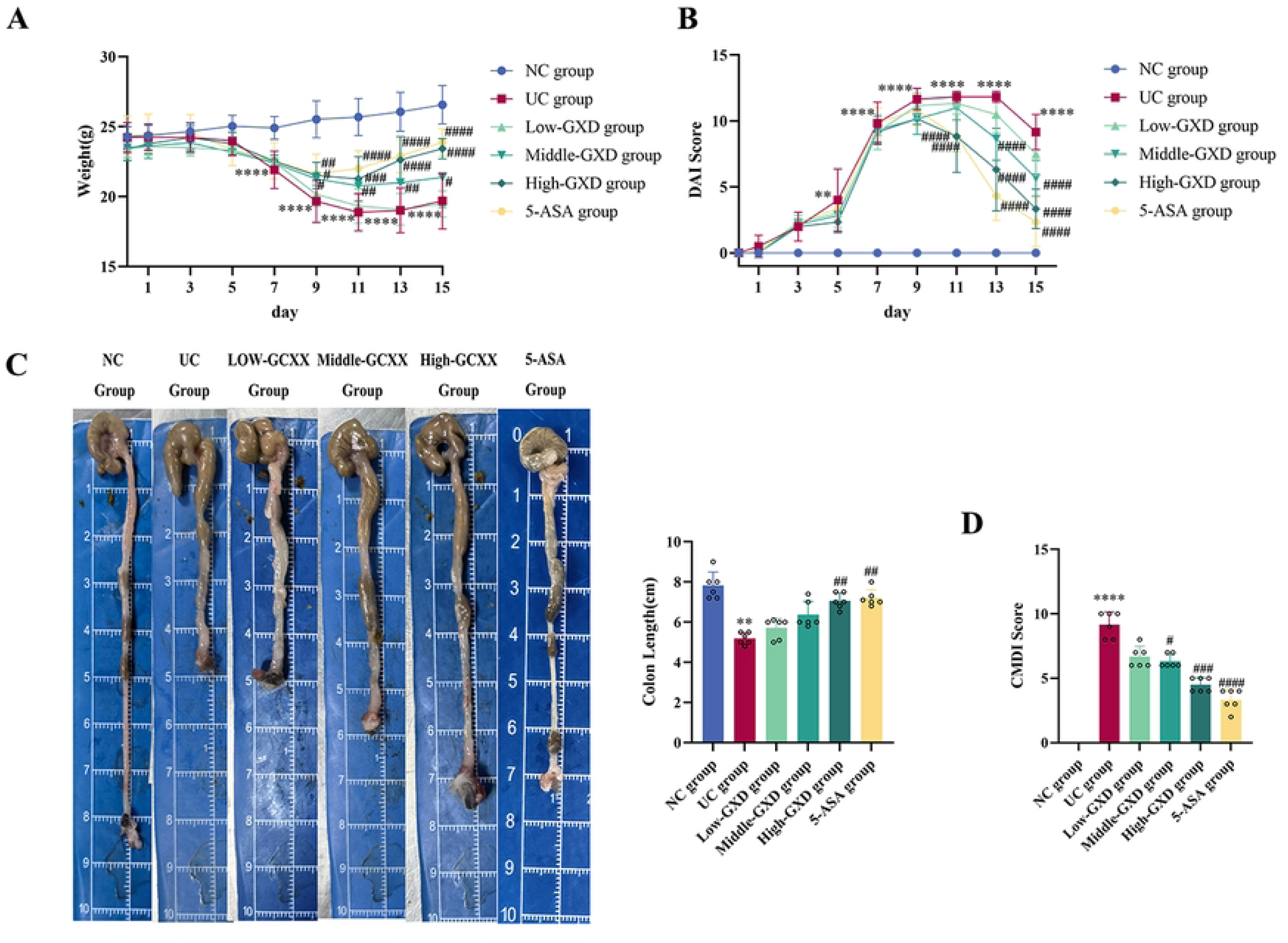
GCXX attenuated colitis in DSS-induced UC. (A) Change in body weight. (B) Change in DAI score during the experiment. (C) Length of colon. (D) Score of colon mucosa injury (CMDI). Compared with the control group **** P <0.0001, *** P <0.001, ** P <0.01, ** P <0.05; GCXX group compared with the model group,####P <0.0001,###P <0.001,##P <0.01,#P <0.05.

### 3.4 GCXX improved colonic mucosal damage in DSS-induced UC mice

Colonic tissue in the control group exhibited intact colonic epithelium with few scattered lymphocytes and eosinophils in the lamina propria. The UC group displayed significant ulceration and defects in mucosal epithelium, with more neutrophil infiltration in lamina propria and submucosa, crypts and crypt abscess, as well as lymphocytes and plasma cells at the base of the mucosa. In contrast, low, medium and high-dose GCXX groups exhibited a marked reduction in inflammatory infiltration and tissue damage, and high-dose GCXX groups exhibited a most marked reduction. UC group were observed to have a much higher colon histopathology score in reference to the NC group, while the low, medium and high-dose GCXX groups showed an even lower score (Fig. 5A). The tight junction proteins Occludin, ZO-1, and Claudin-1 were significantly downregulated in the colon epithelium of DSS colitis mice, whereas their protein expression level was significantly restored after GCXX treatment (Fig. 5B). The UC group presented with significantly higher serum concentrations of DAO and D-Lactate, which in the contrast was observed to be significantly reduced in the low, medium and high-dose GCXX groups (Fig. 5C and D).

**Figure 5.**
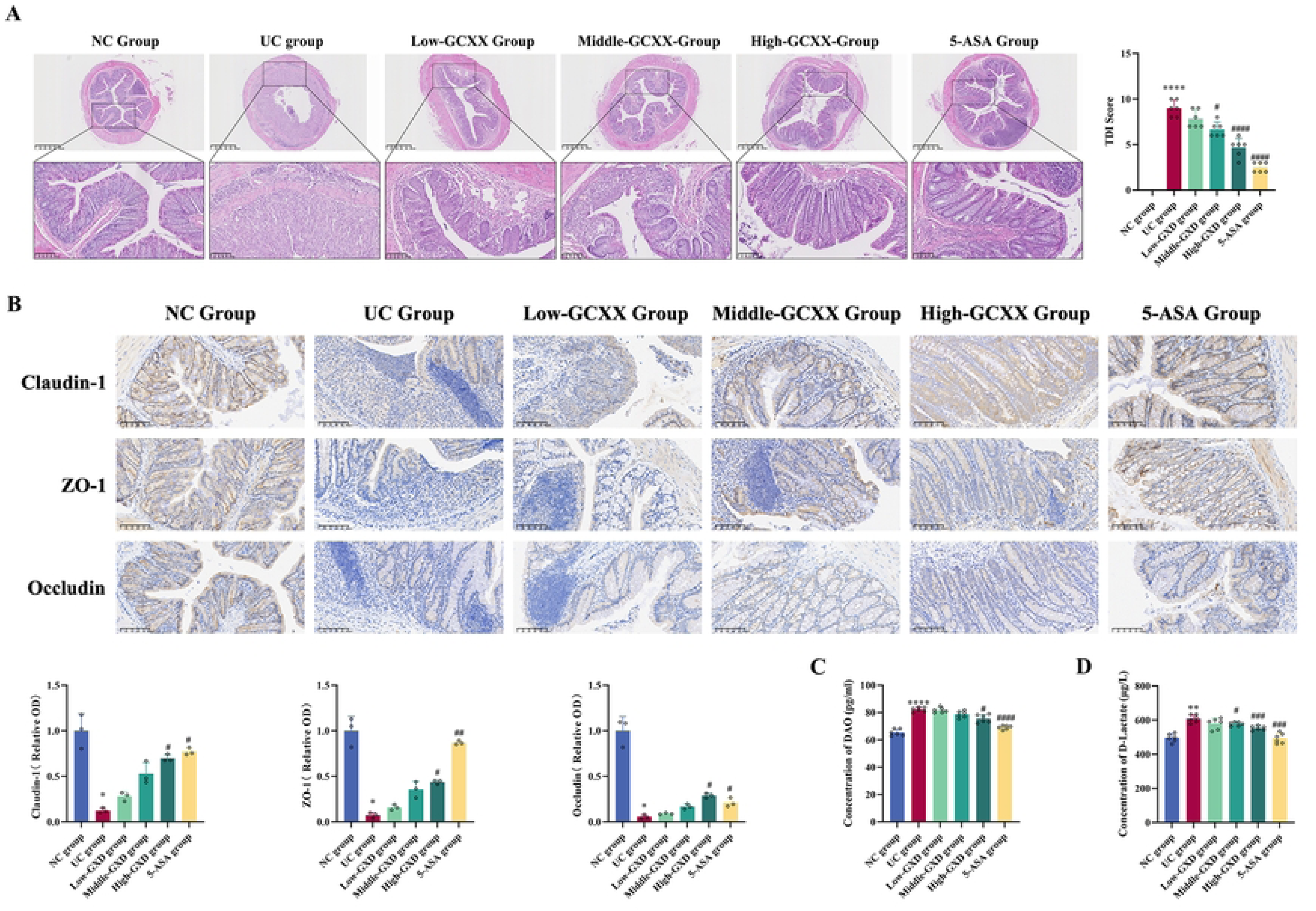
GCXX improved colonic mucosal damage in DSS-induced UC mice. (A) Typical histopathological findings and TDI scores(X5, X20). (B) Distribution characteristics and Occuldin, ZO-1 and Claudin-1 levels (X20). (C) Serum DAO concentration. (D) D-Lactate concentration. Compared with the control group **** P <0.0001, *** P <0.001, ** P <0.01, ** P <0.05; GCXX group compared with the model group,####P <0.0001,###P <0.001,##P <0.01,#P <0.05.

### 3.5 GCXX alleviated OS levels in colon tissues of DSS-induced UC mice

In reference to the control, the MDA was significantly higher in the UC group and decreased in the low, medium and high-dose GCXX groups (Fig. 6A). Additionally, SOD vitality was obviously lower in the colon of UC group and increased in that of low, medium and high-dose GCXX groups (Fig. 6B).

**Figure 6.**
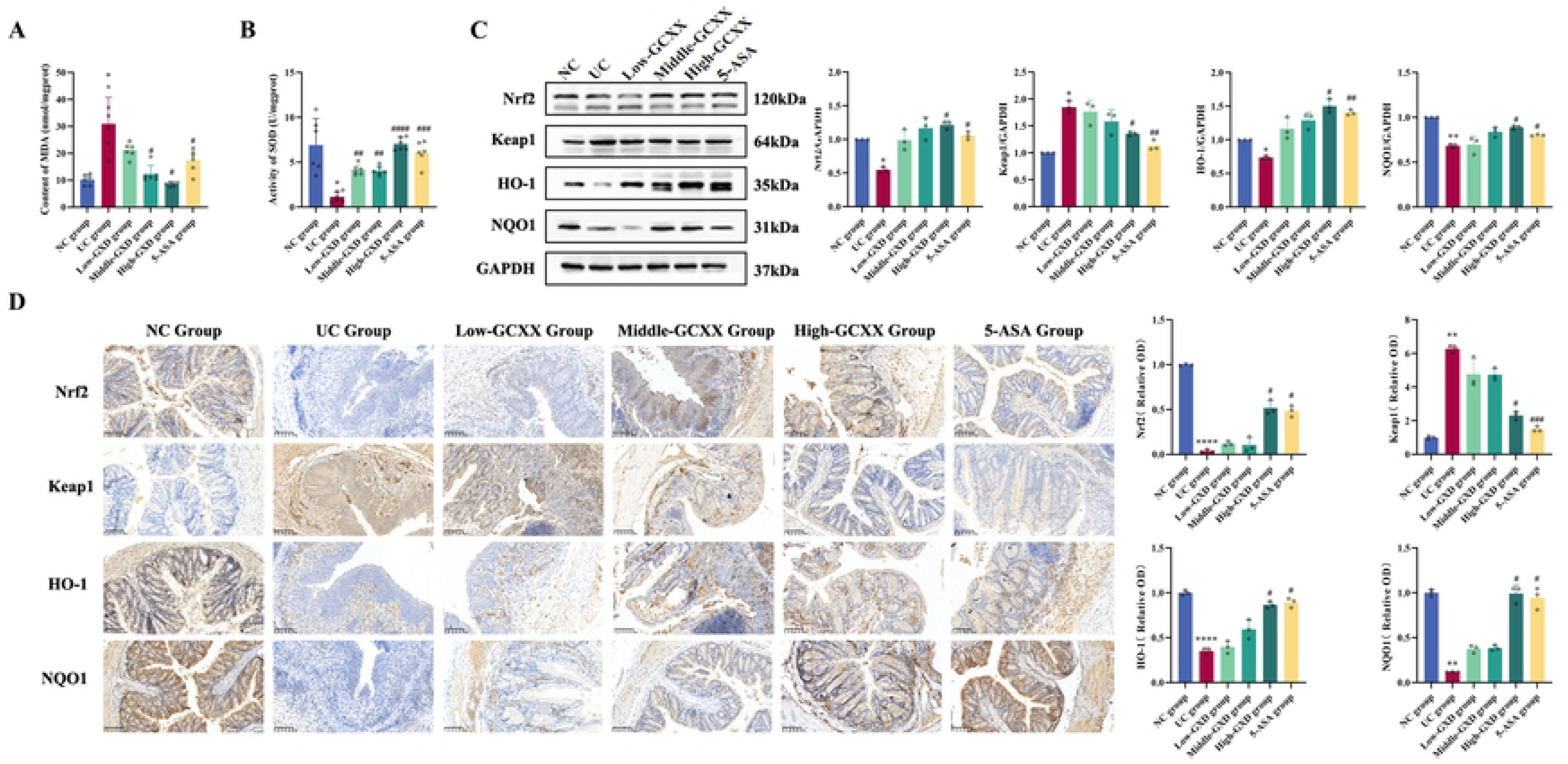
MDA, SOD, and Nrf2 / ARE signaling axis-related proteins expression in mouse colon tissue. (A) MDA concentration. (B) SOD enzyme viability. (C) WB, distribution of Nrf2, Keap1, HO-1 and NQO1. (D) Expression of Nrf2, Keap1, HO-1 and NQO1 in tissues (X20). Compared with the control group **** P <0.0001, *** P <0.001, ** P <0.01, ** P <0.05; GCXX group compared with the model group,####P <0.0001,###P <0.001,##P <0.01,#P <0.05.

### 3.6 GCXX regulated protein expression associated with the Nrf 2 / ARE signaling axis in colonic tissues of DSS-induced UC mice

NQO1, Nrf2 and HO-1 levels in colonic epithelium of low, medium and high-dose GCXX groups mice increased markedly compared to those in UC group; Keap1 protein level notably decreased in the low, medium and high-dose GCXX groups in reference to the UC group, whereas the UC group demonstrated an obvious difference in comparison to the NC group (Fig. 6C and D).

### 3.7 GCXX effectively improved the survival of cells in OS and reduced OS and cell apoptosis

In relation to the control, cell survival was markedly lower in the OS group. Furthermore, notably higher cell survivals were found in both the Low- and High-GCXX groups in contrast to the OS group (Fig. 7A). Compared to the NC group, the OS group showed a marked decrease in SOD enzyme activity and antioxidant capacity, while both parameters were significantly elevated in the Low- and High-GCXX groups (Fig. 7B and C). Similarly, MDA content and apoptosis rate were substantially higher in the OS group, and decreased in the Low- and High-GCXX groups (Fig. 7D). The apoptotic rate and expression of pro-apoptotic protein Bax were obviously elevated, whereas the expression of anti-apoptotic protein Bcl-2 decreased notably in the OS group compared to the NC groups. The apoptotic rate and expression of pro-apoptotic protein Bax were significantly reduced, whereas the expression of anti-apoptotic protein Bcl-2 was significantly higher in the Low-GCXX and High-GCXX groups than the OS groups (Fig. 7E and F).

**Figure 7.**
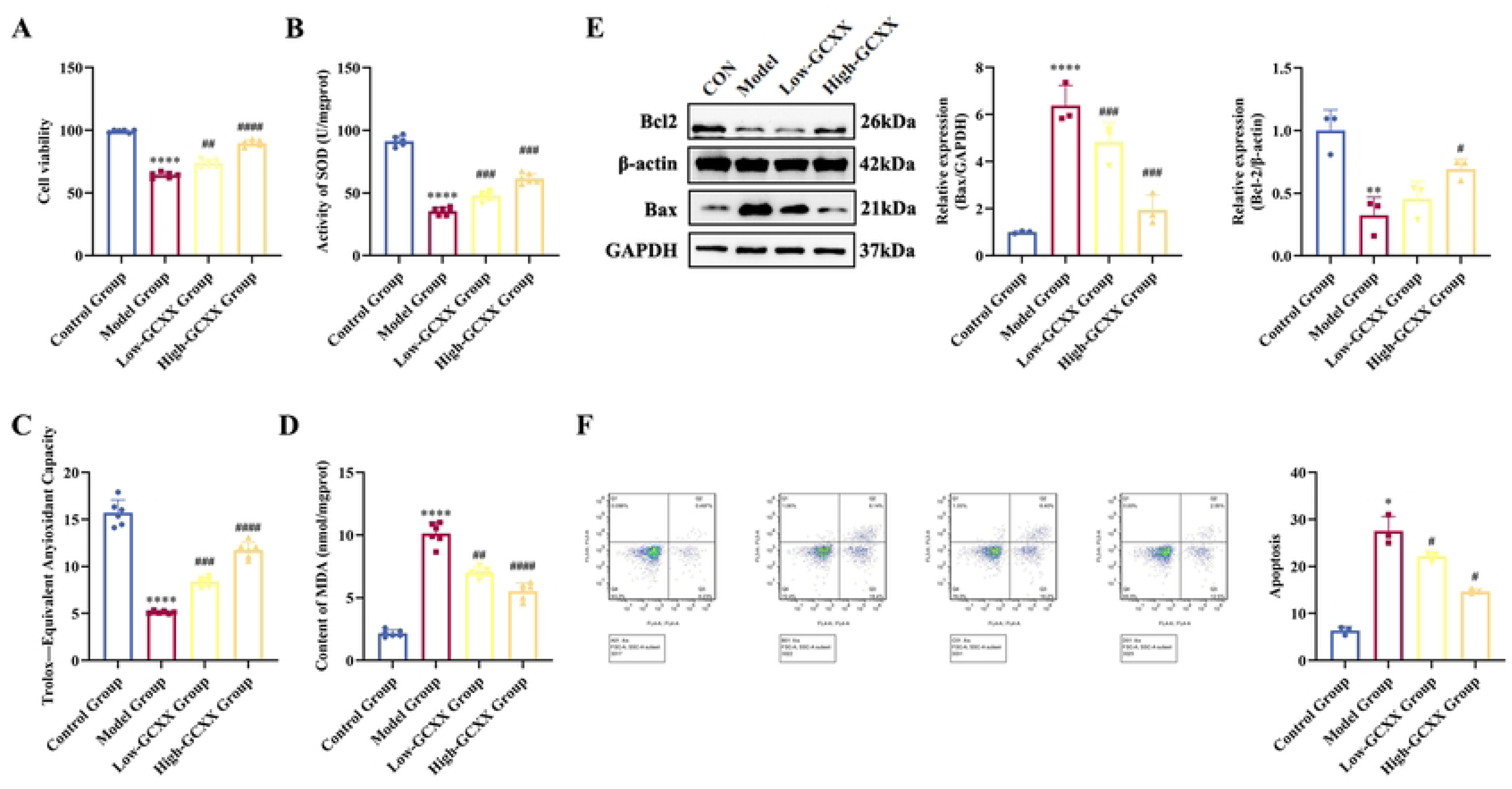
Detection of cell survival, apoptosis and OS. (A) Cell viability by MTT. (B) SOD enzyme viability of cells. (C) MDA content. (D) Total antioxidant capacity of cells. (E) Development and analysis of apoptotic protein. (F) Apoptotic rate of cells. Compared with the control group **** P <0.0001, *** P <0.001, ** P <0.01, ** P <0.05; GCXX group compared with the model group,####P <0.0001,###P <0.001,##P <0.01,#P <0.05.

### 3.8 GCXX improved OS-induced damage in UC by regulating the NRF 2-ARE signaling axis

In comparison to the control, the OS group showed an increase in the relative fluorescence intensity of Nrf2, HO-1, and NQO1, which further. rebounded in the Low- and High-GCXX groups. In addition, further reduced relative fluorescence intensity of these proteins were found in the si-Nrf2 + OS + GCXX group (Fig. 8A-C).

**Figure 8.**
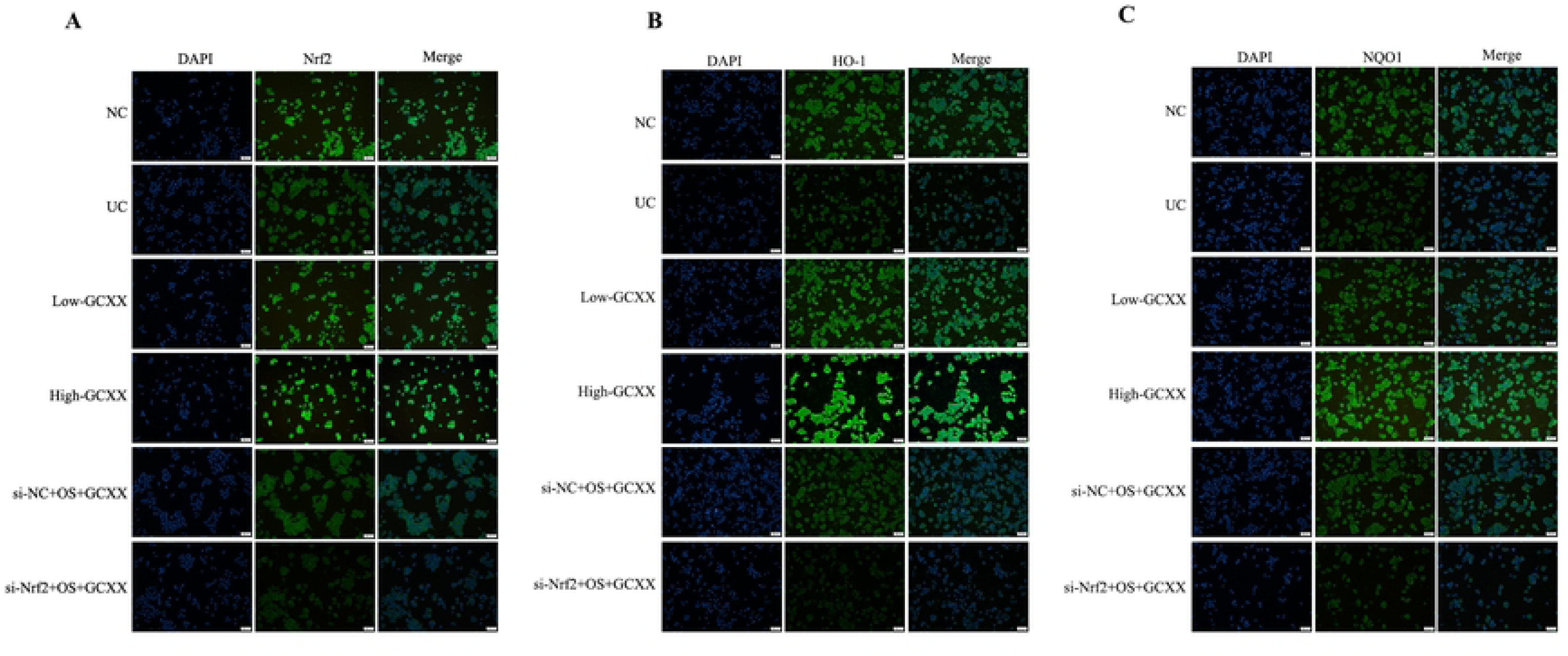
Expression and distribution of key proteins in the NRF2 / ARE signaling axis. (A) Expression and distribution of Nrf2. (B) Expression and distribution of HO-1. (C) Expression and distribution of NQO1.

## 4. Discussion

Traditional Chinese medicine, a treasure handed down over millenni, has unique advantages in UC treatment^[(18)]^. GCXX, a traditional Chinese medicine for ulcerative colitis, has been extensively studied. Studies have shown that GCXX combined with mesalazine yields superior outcomes compared to mesalazine monotherapy ^[(19)]^. In the UC mouse model, GCXX treatment notably alleviated colitis by reguating gut microbiota and their metabolites ^[(8)]^, downregulated the levels of proinflammatory factors like TNF-α and IL-6, thereby ameliorating DSS-triggered colitis. In our current study, the weight, DAI score, colon length and CMDI score of UC mice were significantly recovered after GCXX treatment.

The therapeutic mechanism of GCXX for UC is characterized by multiple pathways, targets and mechanisms. Therefore, we explored the mechanism of GCXX treatment for UC with the help of network pharmacology methods and bioinformatics analysis, and noticed that GCXX may intervene in UC through OS-related signaling pathways. UHPLC-MS/MS results, such as curcumin, baicalin, berberine and other monomer components with clear antioxidant functions ^[(20–22)]^. Berberine can reduce OS-induced damage by modulating the Keap1 / Nrf2 / NF-kappaB signaling axis to mitigate TNBS-induced colitis in rats ^[(12)]^. Moreover, curcumin attenuated OS-caused damage by activating the antioxidant system through activation of mouse macrophages ^[(23)]^.

OS-induced damage results from a pathophysiological process wherein cells produce excessive reactive oxygen species (ROS) and reactive nitrogen species (RNS) in response to various harmful stimuli. ROS and RNS overproduction ultimately causes significant damage to cellular tissues ^[(24)]^. The damaging effects of ROS include activation of inflammatory mediators like NF-kappaB, which mediates the production of many inflammatory cytokines such as TNF-α, thereby eliciting a widespread inflammatory response^[(24–26)]^. Additionally, ROS can destroy the physiological structure and normal function of the host cell through the biofilm of lipid peroxidation cells, or MDA, the lipid peroxidation metabolites of ROS, and inactivate a variety of hormones and enzymes in the body, thereby changing the oxidation state of the host cell and reducing its antioxidant capacity, so that the OS-induced damage will be continuously superimposed ^[(27, 28)]^. Studies have shown that persistent low-grade inflammation can induce OS-induced damage in the colon in the pathogenesis of UC ^[(13)]^, destroy the intestinal mucosal barrier, enhance permeability, and exacerbate local lesions, thereby leading to deteriorated condition. It is interesting to note that the endogenous apoptosis pathway is triggered when there are DNA damage, hypoxia as well as stress states. OS has been demonstrated as one of the key triggers of the cellular endogenous apoptosis pathway in a variety of diseases ^[(29, 30)]^. Similarly, the accelerated apoptosis of intestinal epithelial cells has a significant effect on the intestinal mucosal injury of UC ^[(31, 32)]^. Our study showed that MDA, ROS and apoptosis in enterocytes of OS state increased significantly, while treatment with GCXX could effectively lower MDA, ROS and apoptosis levels in a dose-dependent manner.

The NRF 2 / ARE signaling axis is the clear antioxidant mechanism ^[(33, 34)]^. It acts as a critical pathway within the cellular antioxidant system, and its regulated downstream phase metabolic enzymes and antioxidant proteases play a pivotal role in cellular defense protection ^[(35)]^. Nuclear factor-erythroid 2-related factor 2 (Nrf2), a redox core factor containing seven domains from Neh1 to Neh7 ^[(36)]^, triggers the expression of intracellular phase detoxification enzymes and antioxidant enzymes, improves the body’s antioxidant capacity, promotes cell survival, and maintains the homeostasis of cell oxidation-antioxidant balance. The Nrf2 protein is expressed in the colon tissue ^[(37)]^. Under physiological conditions, Keap1 is a substrate of a dependent E3 ubiquitin ligase complex that promotes ubiquitination of Nrf2 and rapid degradation ^[(38, 39)]^. When exposed to oxidative stress, Nrf2 separates from Keap1, moves into the nucleus, joins tiny Maf proteins in a heterodimer, and binds to the antioxidant response elements (ARE). This binding initiates the transcription and expression of phase II detoxification enzymes and antioxidant proteins, thereby enhancing cellular defense against ROS and improving the antioxidant capacity of local tissues. Studies have confirmed the inverse correlation between Nrf2 and ROS levels in wild-type mice by combining real-time electron paramagnetic resonance imaging and Nrf2 (- / -) mice ^[(40)]^. Superoxide dismutase (SOD) is an important antioxidant enzyme for removing superoxide anion free radicals ^[(41)]^. In one study, the SOD activity of wild-type fibroblasts after treatment with inducer 3H-1,2-dithiole-3-thione (D3T) was markedly elevated, while SOD was not altered in Nrf 2 (- / -) cells, which indicated Nrf2’s role as a key regulator in enhancing SOD’s antioxidant bioactivity in cardiac fibroblasts ^[(42)]^. Heme oxygenase1 (HO-1) is a crucial antioxidant enzyme ^[(43)]^, and HO-1 protein expression is upregulated in response to OS and cellular damage ^[(44)]^. NAD(P)H quinone dehydrogenase 1 (NQO1) is a homodimeric flavinase that catalyzes the reduction of quinone to hydroquinone, thus promoting the excretion of quinone and avoiding ROS production through single-electron reduction reaction and redox cycle (Jung and Kwak, 2010).

Our findings reveal that the level of antioxidant enzyme system represented by SOD is suppressed in OS state, which leads to further imbalance of oxidative-antioxidant balance, continuous increase of ROS and MDA levels, significant decrease of SOD enzyme activity, and continuous aggravation of OS-induced damage. Animal experiments showed that GCXX can effectively elevate Nrf2 protein levels. Compared with the UC group, intestinal barrier function related proteins such as ZO-1,Claudin-1, Occudin and antioxidant proteins such as HO-1 and NQO1 were notably elevated in the GCXX group. However, Nrf2, HO-1, and NQO1 were not obviously different between NC and UC groups, which may be attributed to the complex *in vivo* environment of animals. Because these indicators are regulated by multiple mechanisms, including OS, the final difference between NC and UC groups was not significant. Furthermore, data from available studies suggest that Keap1 has an opposite trend in OS-induced damage ^[(12, 45)]^. The expression of Keap1 was lowered significantly in the UC group, while it further decreased after GCXX administration.

We speculate that in the OS-induced damage state, the body’s defense system may be activated, accelerating the degradation or transcriptional regulation of Keap1 through mechanisms that remain unclear, thus reducing its interaction with Nrf2 and ensuring an adequate amount of Nrf2 for effective antioxidant function. The underlying biological mechanisms warrant further investigation through additional experimental studies. The results of cell experiments showed that GCXX can improve OS-induced damage in UC, activate the antioxidant system, fight OS, improve intestinal epithelial cell survival, and strengthen the intestinal mucosal barrier through the Nrf 2 / ARE signaling axis.

## 5. Conclusions

Here, We prove that GCXX activates the expression of downstream antioxidant proteins by mediating the NRF 2-ARE signaling axis, strengthens the antioxidant system to reduce OS-induced damage in UC, downregulates enterocyte apoptosis, and protects the intestinal mucosal barrier. This study confirms the pharmacological activity of GCXX and provides theoretical support for its clinical application.

## Abbreviations

GCXX: GanCaoXieXin
UC: Ulcerative colitis
UHPLC-MS/MS: Ultra-performance liquid chromatography combined with quadrupole time-of-flight mass spectrometry
DSS: Dextran Sulfate Sodium Salt
IHC: Immunohistochemistry
WB: Western blot
ELISA: Enzyme-linked immunosorbent assay
IF: Immunofluorescence
OS: Oxidative stress
IBD: Inflammatory bowel disease
DAI: Disease activity index
CMDI: Colon macroscopic damage index
TDI: Tissue damage index
SPF: Specific pathogen-free
SD: Sprague-Dawley
TCMSP: Traditional Chinese Medicine Systems Pharmacology Database and Analysis Platform
GO: Gene Ontology
KEGG: Kyoto Encyclopedia of Genes and Genomes
DEGs: Differentially expressed genes
siRNA: Small interfering RNA
SRC: Non-receptor tyrosine kinase
PIK3R1: Phosphatidylinositol 3-kinase
MAPK3: Mitogen-activated protein kinase 3
ROS: Reactive oxygen species
RNS: Reactive nitrogen species
Nrf2: Nuclear factor-erythroid 2-related factor 2
ARE: Antioxidant response elements
SOD: Superoxide dismutase
D3T: 3H-1,2-dithiole-3-thione
HO-1: Heme oxygenase1
NQO1: NAD(P)H quinone dehydrogenase 1

## Ethics approval and informed consent

All animal experiments were approved by the Experimental Animal Ethics Committee of ZCMU (IACUC-20231120-28) and conducted at ZCMU. Animal welfare work was conducted as per the Guidelines on the Management and Use of Laboratory Animals (Ministry of Science and Technology, China, 2016).

## Consent for publication

We confirm that the details of any images, videos, recordings, etc can be published, and that the person(s) providing consent have been shown the article contents to be published.

## Data availability

The data that support the findings of this study are openly available in the GEO database at http://www.ncbi.nlm.nih.gov/geo, reference number[GSE243625].

## CRediT authorship contribution statement

**Ping Ling**: Conceptualization, Writing – original draft. **Bing Zhang**: Validation. **XinRui Zhang**: Investigation. **YingChao Liu**: Investigation, Formal analysis. **Yan Shen**: Writing – review & editing.

